# Molecular Mechanisms of Gain-of-Function Mutations in λ Cro Revealed by Molecular Dynamics Simulations

**DOI:** 10.1101/2025.08.14.670369

**Authors:** Ryan Hebert, Alexander Perez, Jeff Wereszczynski

## Abstract

Transcription factors regulate gene expression by coordinating complex networks in organisms ranging from bacteriophages to humans. Bacteriophage λ Cro is a 66-residue repressor that binds DNA as a dimer to block transcription. Because of its small size, simple structure, and well-characterized function, Cro has long served as a model system for understanding the structure/function relationship in transcription factors. Experiments have shown that a small set of mutations can convert it into a dual-function transcription factor capable of both repression and activation. One engineered variant retains activity when truncated to 63 amino acids but loses function at 59, highlighting how little sequence is required for complex regulatory behavior. To probe the molecular basis of this adaptability, we performed multi-microsecond all-atom molecular dynamics simulations of wild-type Cro and two engineered variants, Act3 and Act8. The simulations reveal that minimal sequence changes can reorganize interaction surfaces, shift DNA-binding modes, modulate binding affinities, and redistribute intramolecular communication pathways. These effects on DNA binding occur alongside changes that may broaden regulatory potential, offering insight into how compact transcription factors evolve new functions. Together, these observations provide a mechanistic framework for understanding how transcription factor sequence, structure, and dynamics reshape gene regulatory function.

## 1 Introduction

Transcription factors play a central role in gene regulation, modulating transcription by altering DNA accessibility both locally and at distal sites.^1–4^ They bind directly to DNA, often recognizing specific sequences, and depending on context and conformation can function as activators, repressors, or dual regulators that do both.^3,4^ Dual-action transcription factors are particularly interesting because they integrate activation and repression in a single protein, but most examples are relatively large, often forming higher-order oligomers or scaffolds that participate in large-scale DNA remodeling.^5–7^ In contrast, some functional repressors are much smaller and structurally simpler, ^8,9^ retaining only the core elements required for DNA binding and regulation. These minimal repressors provide a tractable framework for probing the structural basis of transcriptional control.

One well-studied transcription factor is λ Cro, a 66-residue protein that functions as a compact transcriptional repressor and contains the core elements required for DNA binding and regulation.^10,11^ It has been extensively used as a model system for understanding transcription factor behavior, in part because of its small size and simple, well-resolved architecture. Its role in the λ genetic switch^12,13^ has made it a central example in studies of gene regulatory feedback.^14^ Cro has been used to probe multiple aspects of DNA recognition, from nonspecific recruitment along DNA^15,16^ to high-affinity binding at specific operator sequences.^17,18^ Its helix–turn–helix DNA-binding domain resembles that of the larger λ cI repressor,^19^ yet the overall protein is far smaller, making it an attractive target for engineering additional functionality or altering its properties.^20,21^

Early work by Bushman *et al.* attempted to convert Cro into a transcriptional activator, demonstrating that even small repressors can be modified toward dual regulatory roles.^22^ More recently, Brödel *et al.* used accelerated evolution to create an array of mutant Cro variants.^23^ Using optical density measurements of mCherry and green fluorescent protein (GFP) expression on either side of a promoter region, they identified variants that functioned as dual-action transcription factors. These variants, labeled Act1 through Act17, contained mutations at residues 17, 21, 22, 26, and 31 (Figure 1). As few as three mutations and as little as 63 amino acids in length were sufficient to gain this function typically found in much larger transcription factors and complexes, pushing the limits of what minimal structure is required for gene activation. They also observed a loss of function when these mutants, and in particular Act3, were truncated to 59 amino acids.

**Figure 1:**
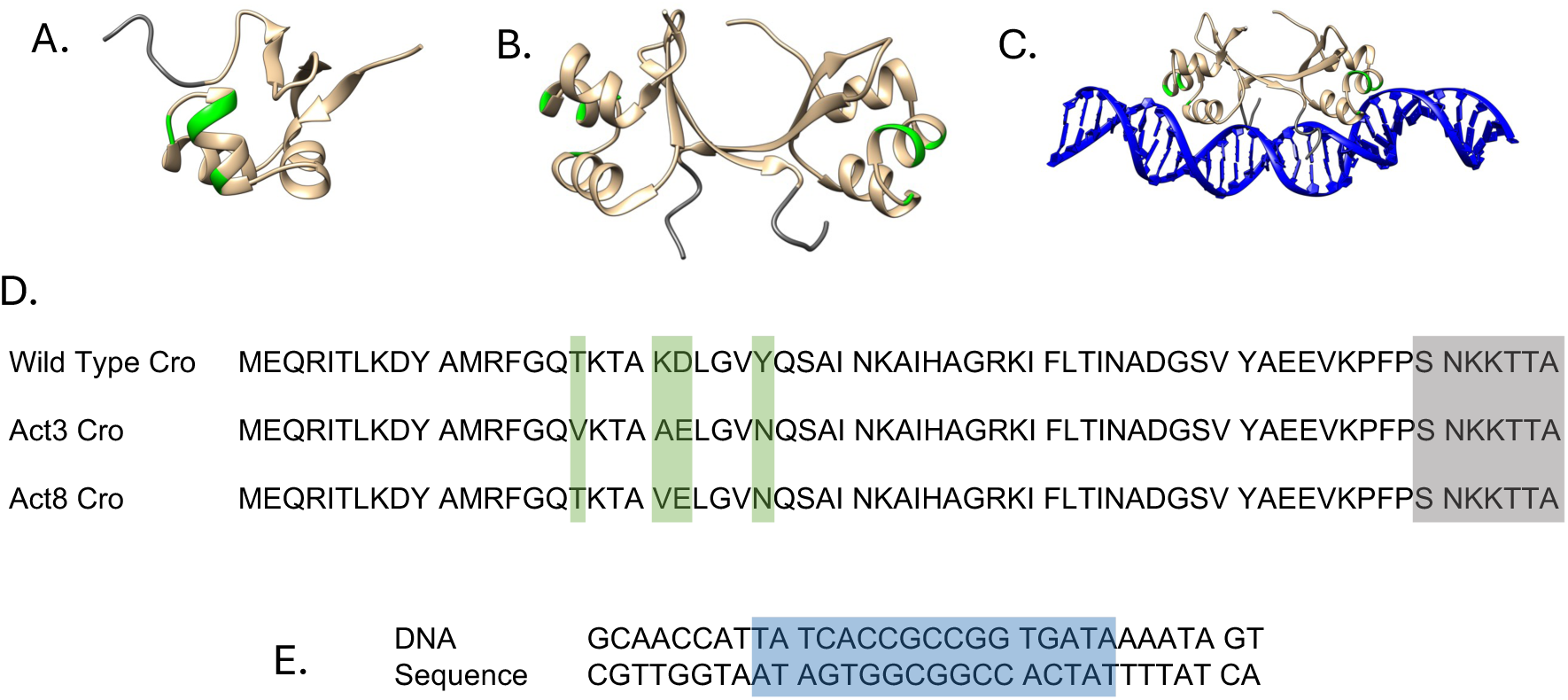
Representations of the Cro systems simulated in this study. Post-equilibration frames are shown for the monomer (A), dimer (B), and DNA-bound complex (C). Mutation sites are highlighted in green, and deletion targets are shown in gray. The sequences of the Cro variants (D) use the same color scheme to indicate mutations and deletions. The DNA sequence (E) highlights the operator region for Cro binding in blue.

Despite these fascinating experimental results, the structural basis for how such mutations convert Cro from a repressor into a dual-function transcription factor remains unknown. While experiments show that a small number of mutations can produce a drastic change in function, they do not reveal how these changes remodel Cro’s structure and dynamics to achieve this transformation. To uncover the structural basis for these functional changes, we performed atomic-scale molecular dynamics (MD) simulations of wild-type Cro and two previously characterized dual-function variants, Act3 and Act8. The simulations reveal that mutations remodel a region of the protein’s surface, forming a distinct interaction patch with altered hydrogen-bonding patterns and surface geometry. Contact network analyses suggest that these changes promote cooperative accessibility of this patch, potentially enhancing interactions with binding partners. The disordered C-terminal region emerges as a dynamic, context-sensitive element that appears to act as a structural probe. Mutations also subtly alter interaction networks in the DNA-bound forms, suggesting compensatory and cooperative changes to Cro’s architecture to support function. While our simulations focus on DNA-bound forms and do not capture potential effects on RNA polymerase recruitment, they establish a mechanistic foundation for distinguishing between mutations that influence direct DNA contacts and those that may act through polymerase interactions.

## 2 Methods

### 2.1 System Construction

Models of Cro were based on the Protein Data Bank (PDB) entries 1COP^24^ and 6CRO.^25^ The 1COP NMR ensemble was used because it contained complete Cro models. Cro monomers were constructed by deleting the dimer partners from the 1COP ensemble and introducing mutations using the Dunbrack 2010 rotamer library in UCSF Chimera,^26,27^ following the Act3 and Act8 variants from the accelerated evolution experiment.^23^ Cro dimers were constructed in the same way, using part of the NMR ensemble as a starting conformation and applying a similar mutation protocol. DNA-bound Cro systems used the biological assembly of the 6CRO crystal structure. Full models of Cro from the 1COP ensemble were aligned to the incomplete crystal structure of Cro, replacing the missing regions. The DNA model in this process was joined and extended to make a total of 32 base pairs, including the operator region. Unbound DNA systems were made by deleting Cro. The DNA and Cro sequences are shown in Figure 1. Truncation of Cro was accomplished by deleting residues 60–66 in each Cro subunit. In total, this gave three different contexts: free monomers, dimers, and dimers bound to DNA. Within these contexts were six different Cro sequences representing the wild type, Act3, and Act8 variants in both full-length and truncated forms.

### 2.2 Molecular dynamics simulations

Solvation and generation of parameter and starting coordinate files were performed in tleap and parmed from AmberTools.^28,29^ All simulations used the ff19SB force field,^30^ OPC water model,^31^ and Li and Merz 12-6 OPC water ions.^32,33^ Simulations with DNA also used the BSC1 DNA force field.^34^ Cro monomers were solvated in water boxes with a 10 Å buffer. Dimer systems were solvated in isometric water boxes with a minimum 10 Å buffer. Crodimers bound to DNA were aligned along an axis and solvated with a 12 Å buffer from the DNA ends and a 20 Å buffer from the sides. All systems were neutralized with Na^+^ and Cl^−^ ions, and additional Na^+^ and Cl^−^ ions were added to give final ion concentrations of 150mM NaCl. To decrease computation time, hydrogen mass repartitioning^35^ was employed in parmed, changing the masses of all non-solvent hydrogens to three AMU and compensating by reducing the mass of the attached heavy atom.

Systems were minimized twice for 10,000 steps, switching from steepest descent to conjugate gradient after 5,000 steps. The first minimization applied a 10 kcal/mol *·*Å^-2^ harmonic restraint to all heavy atoms, and the second minimization had no restraints. Structures were then heated from 5 to 300K over 5ps using a Berendsen thermostat^36^ and Langevin dynamics in an NVT ensemble^37^ with 10 kcal/mol *·*Å^-2^ harmonic restraints on solute heavy atoms. These restraints were progressively reduced by factors of 1/3 every 100 ps in the NPT ensemble and completely removed after 600 ps. All systems after relaxation were simulated in five copies each using pmemd.cuda^38–40^ for up to an additional 5.0 µs in the NPT ensemble at 300K, with 4fs timesteps, a 10 Å non-bonded cutoff, and particle-mesh Ewald for long-range electrostatics^41^ using local resources. Based on root mean square deviation from the first frame (RMSD) data calculated using cpptraj,^28,29^ the first 200ns of each simulation were considered equilibration time for analyses. All minimization, heating, relaxation, and production simulations were performed using Amber24.^28^ Simulations of Cro bound to DNA that featured disassociation of a Cro subunit were stopped after disassociation to prevent periodic image interactions.

### 2.3 Simulation Analyses

Root mean squared fluctuations (RMSF), hydrogen bonding, and solvent accessible surface area (SASA) were measured using cpptraj in AmberTools,^28,29^ while contact analysis was performed with MDAnalysis.^42,43^ Visualization and image processing were done in VMD^44^ and UCSF Chimera.^26^ All time series analyses used all simulation frames, whereas other analyses were carried out on post-equilibration frames only. RMSFs were obtained after RMSD alignment of trajectories to the heavy atoms of residues 7–45 (helices and a β-strand). Average atomic positions of Cro heavy atoms were calculated, and RMSFs for each subunit in each simulation were computed relative to these averages. Values from all trajectories in a system were averaged and are reported as ensemble means with standard errors.

Contacts were defined by a 4.5 Å heavy-atom cutoff, hydrogen bonds by a donor–acceptor cutoff of 3.0 Å and angle of 135°. SASA was calculated in AmberTools using the linear combinations of pairwise overlaps method,^45^ and relative SASA (rSASA) was obtained by dividing each amino acid’s SASA by its empirically derived maximum value in Gly–X–Gly tripeptides.^46^ SASA and rSASA values for mutation targets from both subunits in dimers were combined into a single distribution. MM/GBSA^47^ calculations were performed with MMPBSA.py^48^ in Amber24^28,29^ using the three-trajectory approach: DNA trajectories as receptor, Cro dimers as ligand, and Cro–DNA complexes as the combined system. Post-equilibration frames with contact between both subunits and DNA were used, complex trajectories were run one at a time with igb = 8 (mbondi3 radii set^49,50^), and decorrelation times were determined using pymbar^51,52^ on the complex energy timeseries.

Difference contact network analysis (dCNA) and community detection were performed using publicly available dCNA scripts from Yao and Hamelberg.^53,54^ Average contact probability matrices were generated for each system, and difference maps were obtained by subtracting mutant values from wild type values. The modularity-based algorithm in the same framework was used to identify residue communities within the contact network, enabling detection of shifts in intramolecular communication upon mutation. These analyses were applied to Cro dimers and DNA-bound Cro systems.

## 3 Results

To evaluate how engineered mutations affect Cro’s structural dynamics and functional interactions, we performed molecular dynamics (MD) simulations of wild-type Cro and the Act3 and Act8 variants, which differ at four positions located within or adjacent to the DNA-binding helix-turn-helix motif, including residues in helix 2 (17–22) and the loop preceding helix 3 (residue 26). Each variant was modeled in monomeric, dimeric, and DNA-bound dimeric forms, in both full-length and C-terminally truncated states, yielding 18 systems in total. All-atom simulations were performed in explicit solvent for 5.0 µs across five replicates per system (Table 1).

**Table 1:** Summary of Cro simulations. Each simulation was run in 5 replicates targeting 5 µs per replicate. Actual total time reflects the combined simulation time of all replicates for each system.

| Variant | State | Target Time (μs) | Replicates | Final Time (μs) |
| --- | --- | --- | --- | --- |
| Wild Type (1-66) | Monomer | 5.0 | 5 | 25.0 |
| Wild Type (1-59) | Monomer | 5.0 | 5 | 25.0 |
| Act3 (1-66) | Monomer | 5.0 | 5 | 25.0 |
| Act3 (1-59) | Monomer | 5.0 | 5 | 25.0 |
| Act8 (1-66) | Monomer | 5.0 | 5 | 25.0 |
| Act8 (1-59) | Monomer | 5.0 | 5 | 25.0 |
| Wild Type (1-66) | Dimer | 5.0 | 5 | 25.0 |
| Wild Type (1-59) | Dimer | 5.0 | 5 | 25.0 |
| Act3 (1-66) | Dimer | 5.0 | 5 | 25.0 |
| Act3 (1-59) | Dimer | 5.0 | 5 | 25.0 |
| Act8 (1-66) | Dimer | 5.0 | 5 | 25.0 |
| Act8 (1-59) | Dimer | 5.0 | 5 | 25.0 |
| Wild Type (1-66) | DNA-bound | 5.0 | 5 | 22.7 |
| Wild Type (1-59) | DNA-bound | 5.0 | 5 | 25.0 |
| Act3 (1-66) | DNA-bound | 5.0 | 5 | 21.9 |
| Act3 (1-59) | DNA-bound | 5.0 | 5 | 12.52 |
| Act8 (1-66) | DNA-bound | 5.0 | 5 | 25.0 |
| Act8 (1-59) | DNA-bound | 5.0 | 5 | 25.0 |

### 3.1 Cro Mutations Form a New Interaction Patch

Our MD simulations show that the Act3 and Act8 mutations alter Cro’s surface near helices 2 and 3, with reduced DNA engagement and greater exposure of a chemically distinct patch. To characterize these effects, we examined residue–DNA and residue–protein interactions using both contact frequency and hydrogen bonding analyses. Most interactions involved helices 2 and 3, which flank the mutation sites and are positioned at the DNA interface.

We examined contacts and hydrogen bonds for each of the four mutation sites (residues 17, 21, 22, and 26) to determine how interactions changed in the Act3 and Act8 mutants. Residue 17 consistently contacted DNA in both wild-type and mutant proteins (Figures S1–S3). In DNA-bound wild-type Cro, Thr17 formed two hydrogen bonds with thymine in approximately 85% of frames (Figure 2); in the Act3 mutant, where Thr17 was replaced by valine, only a single hydrogen bond was observed in about 11% of frames. Residue 21 contacted DNA in approximately 20–25% of frames in wild type, but this interaction was eliminated when Lys21 was mutated to alanine or valine. Residue 22 more frequently contacted the protein’s β-sheet than DNA. In wild-type Cro, Asp22 formed hydrogen bonds with Lys18 in about 70% of DNA-bound frames and with Tyr10 in 10–32% of frames; in the mutants, substitution with glutamic acid reduced bonding with Lys18 to roughly 50% of frames and eliminated hydrogen bonding with Tyr10 and other parts of Lys18. Residue 26 consistently contacted DNA in both wild-type and mutant proteins. In wild-type, Tyr26 formed a hydrogen bond with guanine in 67–80% of frames; in the mutants, substitution with asparagine reduced guanine bonding to 43–50% of frames while introducing new hydrogen bonds with thymine in 20–40% of frames.

**Figure 2:**
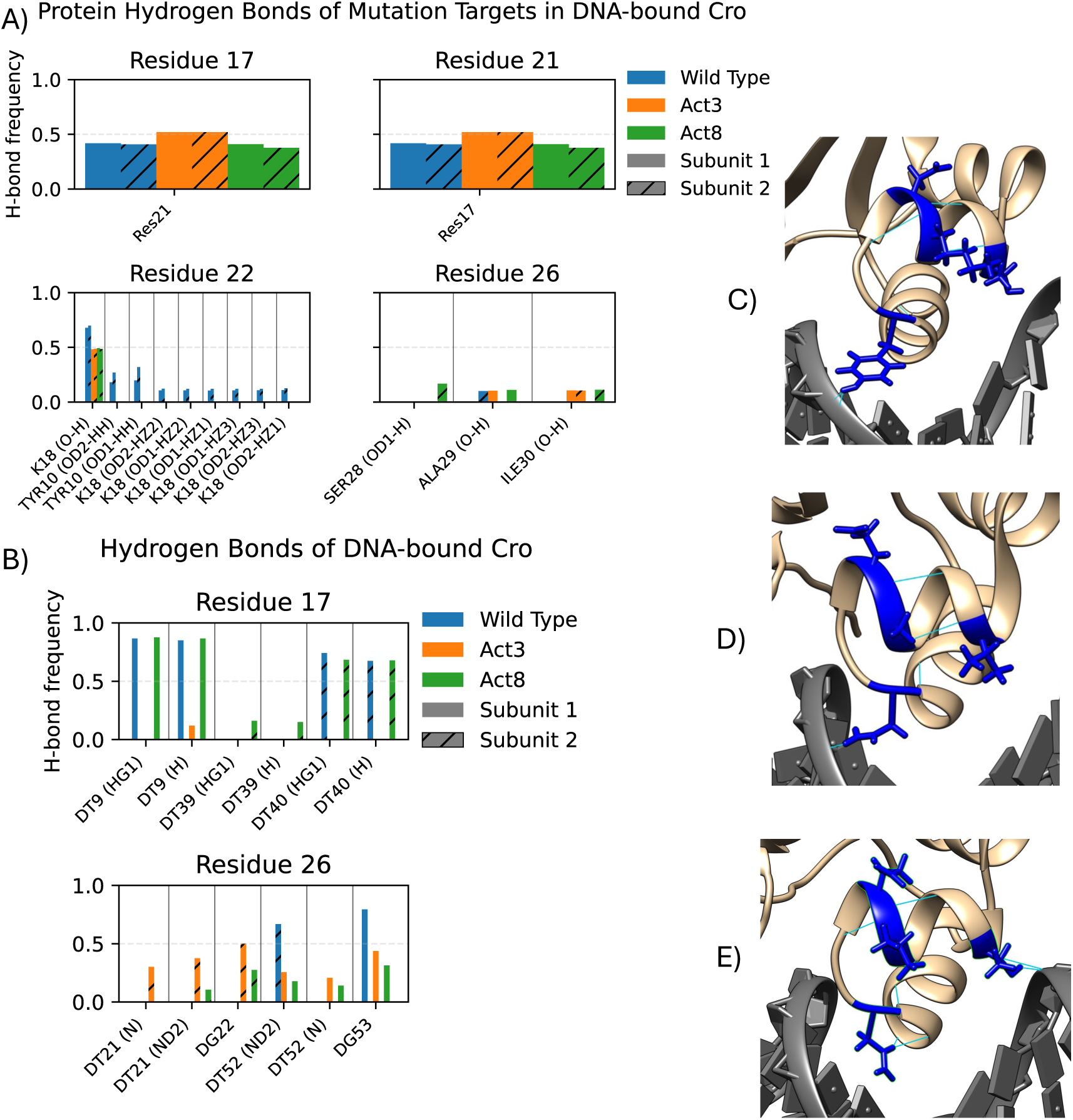
Hydrogen bond analysis of Cro mutation targets in DNA-bound systems. Protein binding partners (A) and DNA binding partners (B) show different average bond formation. These differences are traceable to the mutation from wild type (C) to Act3 (D) and Act8 (E).

We next quantified solvent exposure at the four mutation sites using both solvent accessible surface area (SASA) and relative SASA (rSASA; standard SASA values are provided in Figure S4). We focus on rSASA here, as it normalizes exposure by residue size and avoids conflating side chain volume with accessibility. In our wild-type Cro simulations rSASA values for the four mutated sites ranged from 0.265 to 0.399, indicating moderate burial near the protein surface or at protein–DNA interfaces (Figure 3). Upon mutation, three residues showed increased rSASA: residue 22 increased modestly to 0.425 *±* 0.041, while residues 17 and 21 reached values between 0.567 and 0.717. These shifts reflect greater solvent exposure in the mutants and are consistent with the observed loss of DNA contacts and hydrogen bonds at these positions. The increased exposure of residues 17 and 21, in particular, may make these side chains more available for solvent-mediated interactions or binding to alternative partners.

**Figure 3:**
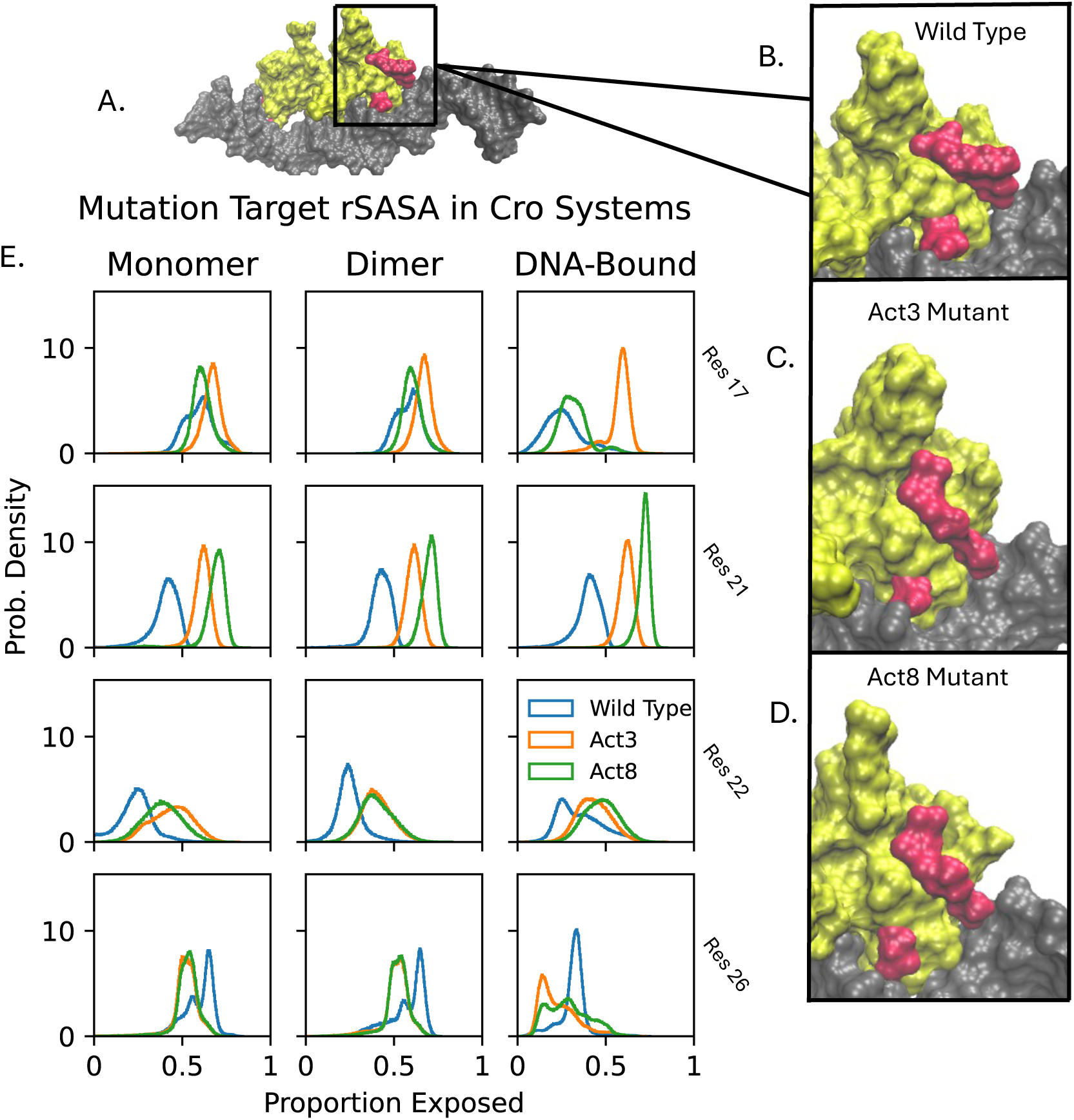
Relative solvent accessible surface area (rSASA) analysis of Cro bound to DNA. DNA is shown in gray, mutation sites in red, and the remainder of the Cro protein in yellow. The overall complex (A) maintains a stable structure, while local remodeling at mutation sites alters the protein surface (B–D). rSASA values (E) indicate changes in burial for each mutation site.

Taken together, the interaction and rSASA results from our simulations suggest that the Act3 and Act8 mutations reduce DNA engagement by eliminating several native protein–DNA and protein–protein contacts, particularly at residues 17, 21, and 22, while increasing solvent exposure at multiple sites. The resulting chemically distinct surface patch may allow these regions to form alternative intermolecular interactions, potentially enabling binding partners other than the original DNA target. These findings indicate that the mutations do more than disrupt existing contacts; they appear to reconfigure local structural networks in a way that could expand Cro’s functional repertoire beyond that of the wild-type protein.

### 3.2 Cro’s Disordered C-terminal Tail Redistributes Interactions Based on Assembly State

We next examined the behavior of Cro’s disordered C-terminal region (residues S60–A66) across wild type, Act3, and Act8. Although this tail lies outside the structured core, its sequence is conserved across variants, and prior experimental work showed that truncating it impairs activity in some mutant contexts. We therefore asked whether this region exhibits persistent flexibility and whether it interacts transiently with other molecular surfaces in a way that might support Cro function.

Across all full-length simulations of wild type, Act3, and Act8, the C-terminal tail was the most flexible region of the protein, with RMSFs substantially higher than those of the structured core (L7–N45), which stayed below 3.4 Å. Tail flexibility was greatest in monomeric simulations, reduced upon dimerization, and further suppressed in DNA-bound complexes. Maximum tail RMSFs ranged from approximately 19–21 Å in monomers, 14–16 Å in dimers, and 11–13 Å in DNA-bound systems (Figures 4, S5–S6). All three variants followed this trend, with no major differences in tail dynamics under equivalent conditions.

**Figure 4:**
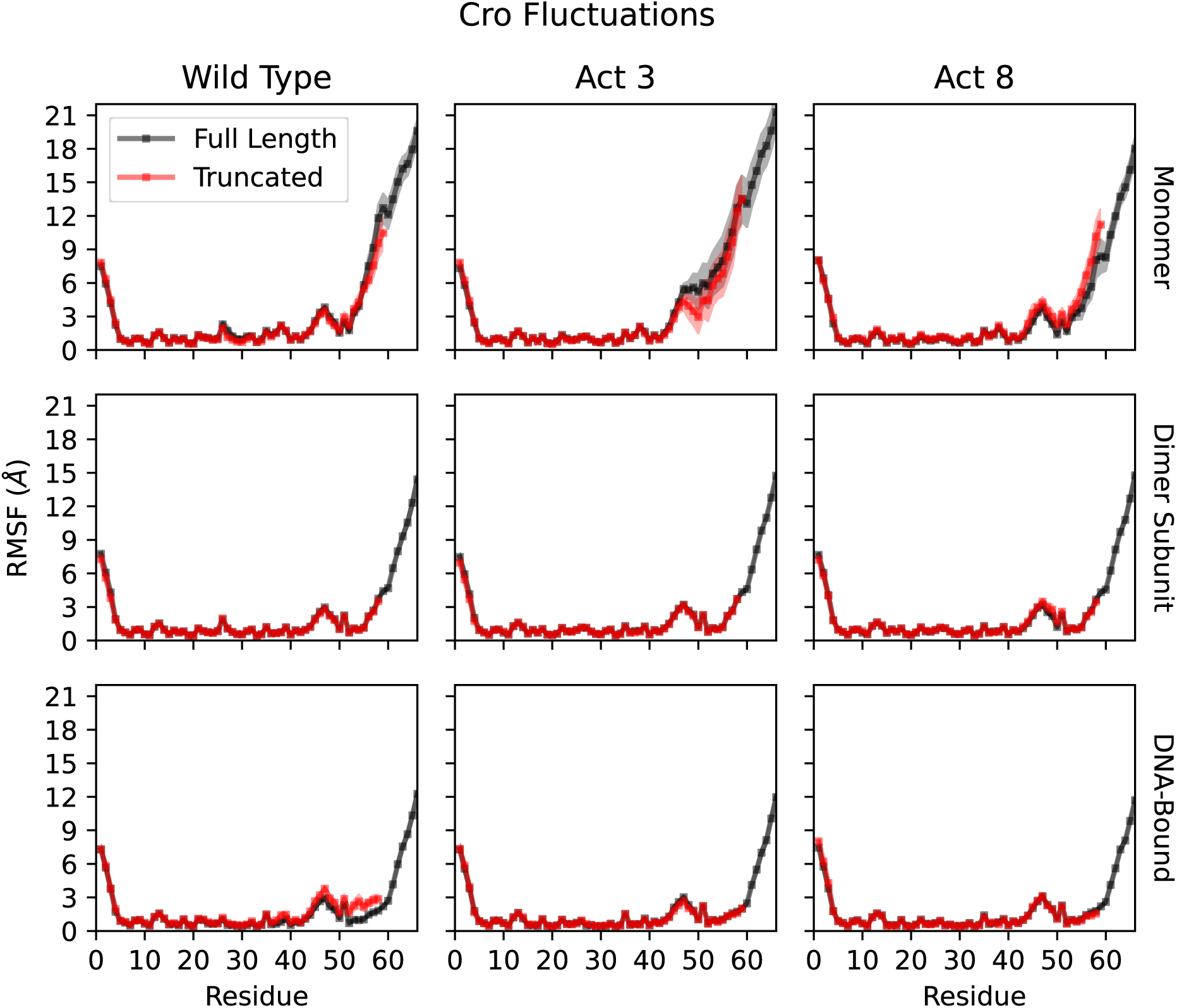
Root mean squared fluctuations (RMSFs) of a single Cro subunit. Top: Monomer fluctuations demonstrate flexibility of the N and C termini, including increases due to unfolding of Cro’s final β-strand. Middle: Dimerized Cro experiences stabilized conformation states while the C-terminus remains flexible. Bottom: Binding to DNA further reduces fluctuations from the average structure while C-terminus fluctuations remain high.

This flexibility allowed the C-terminal tail to dynamically shift its interaction partners according to Cro’s structural state. In monomer simulations, tail–core interactions occurred in approximately 50% of post-equilibration frames, primarily involving contacts with the same subunit’s folded core (residues 1–50) (Figures 5, S7–S9). Upon dimerization, contacts with the same subunit’s core dropped to around 10%, while new interactions with the opposing subunit’s core appeared in roughly 50% of frames, indicating a shift toward inter-subunit stabilization. In DNA-bound systems, a further transition occurred: tail–DNA interactions were observed in over 90% of frames for all three variants, highlighting a dominant role in nucleic acid engagement. Tail–core contacts in this state were more modest, occurring 12–20% of the time with the same subunit and 30–40% with the opposing core for wild type and Act8, whereas Act3 showed somewhat higher opposing-core interactions, reaching 54% of frames in both subunits.

**Figure 5:**
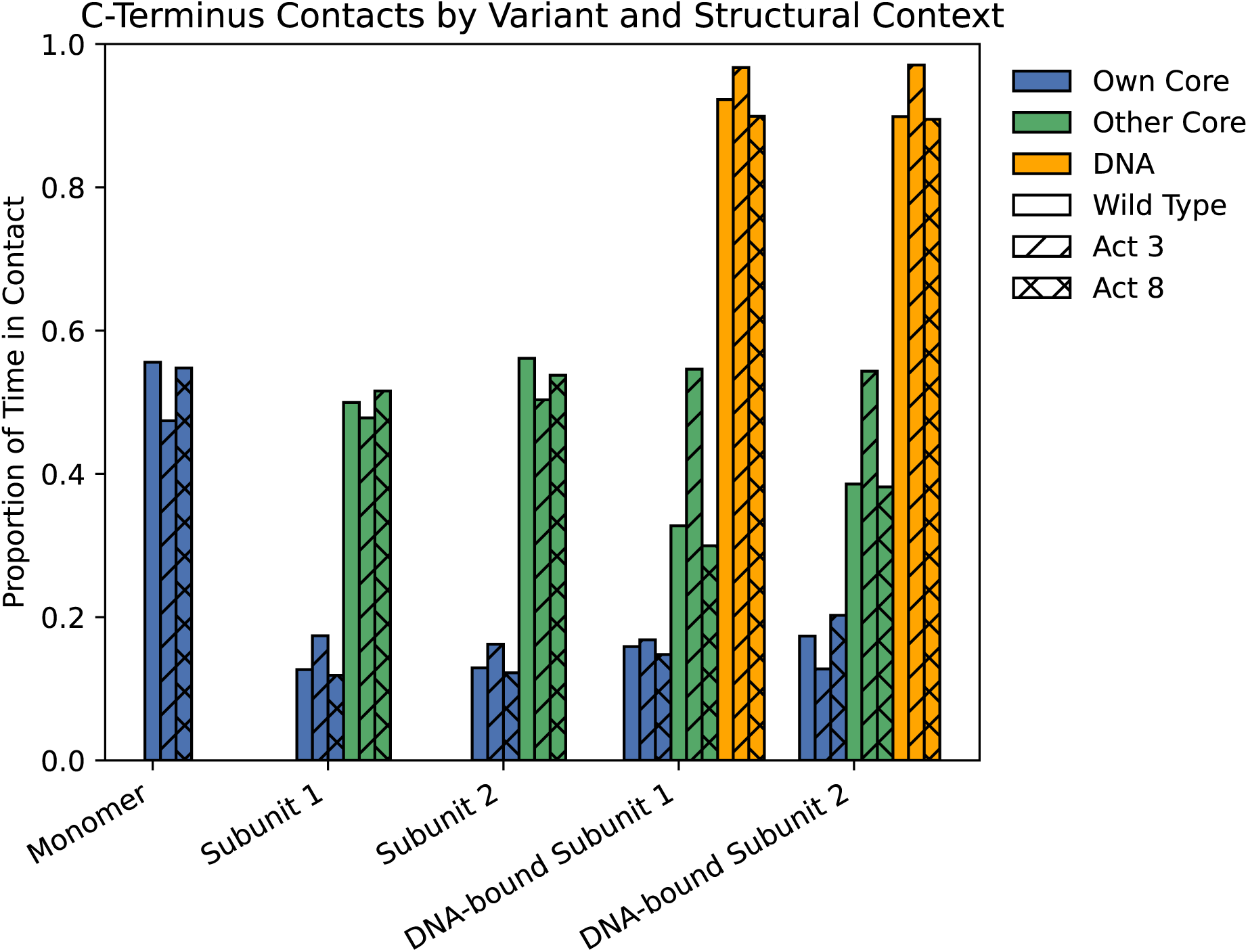
Proportions of simulation time in which Cro’s C-terminus residues 60-66 made contact with other portions of the system.

Overall, these findings indicate that Cro’s C-terminal tail functions as a flexible, disordered probe that dynamically shifts its contacts based on structural context, moving from intramolecular in monomers, to intermolecular in dimers, to persistent DNA engagement in complexes. Its mobility allows it to scan neighboring surfaces and form transient stabilizing interactions, with tail–DNA contacts occurring in over 90% of frames across all variants. The consistent behavior across variants suggests a conserved role for the tail, while the higher opposing-core contact frequency in Act3 DNA-bound systems points to a subtle variant-specific difference in core–tail engagement. Given this behavior, we next asked whether removing the tail alters Cro’s structural stability or its ability to bind DNA.

### 3.3 C-terminal truncation reduces DNA binding in Act3 Cro

Prior experimental work showed that truncating Act3 to 59 amino acids eliminates its function.^23^ Because our analysis in the previous section revealed frequent contacts between the truncated residues and other parts of the system, we asked whether their removal would affect Cro’s structural stability or its ability to bind DNA. Although we observed frequent contacts between these residues and other parts of the system, their loss did not appear to weaken Cro’s dimer structure. No significant difference in fluctuations was observed along the length of each Cro subunit as a result of truncation, suggesting that the β-sheet structure at the interface was sufficiently well integrated to maintain dimer association and that truncation alone is not sufficient to break already associated Cro dimers.

Strikingly, the combination of mutation and truncation had a drastic effect on Cro’s DNA-binding ability in our simulations. A truncated Cro Act3 lost association between DNA and one of its subunits within 3.5 µs in 4 out of 5 simulations. This loss of binding was far more frequent than in full-length systems: only 1 out of 5 simulations of full-length wild type or Act3 experienced a similar dissociation, and that occurred within 2.2 µs. Notably, no dissociation events were observed for full-length Act8, truncated Act8, or truncated wild-type Cro, underscoring that this loss of DNA binding is specific to the Act3 mutant (Figure S10).

To pinpoint the cause of this Act3-specific instability, we compared DNA contact patterns and calculated Cro–DNA binding energies for full-length and truncated forms. For wild type and Act8, truncation reduced contacts almost entirely due to loss of the deleted residues: combining results from both subunits, wild-type Cro residues 1–59 maintained 360*±*14 contacts with DNA before and after truncation, while Act8 went from 338*±*19 to 328*±*15 in the same region. Residues 60–66 made 58*±*18 and 67*±*19 contacts for wild type and Act8 Cro, respectively. In contrast, Act3 Cro residues 1–59 dropped from 350*±*20 to 321*±*16 contacts with DNA, in addition to the loss from truncating residues 60–66 (Figure 6). This extra reduction indicates that the disordered C-terminus plays a particularly important role in maintaining DNA association in Act3, whereas Act8 can sustain stable DNA binding even without it.

**Figure 6:**
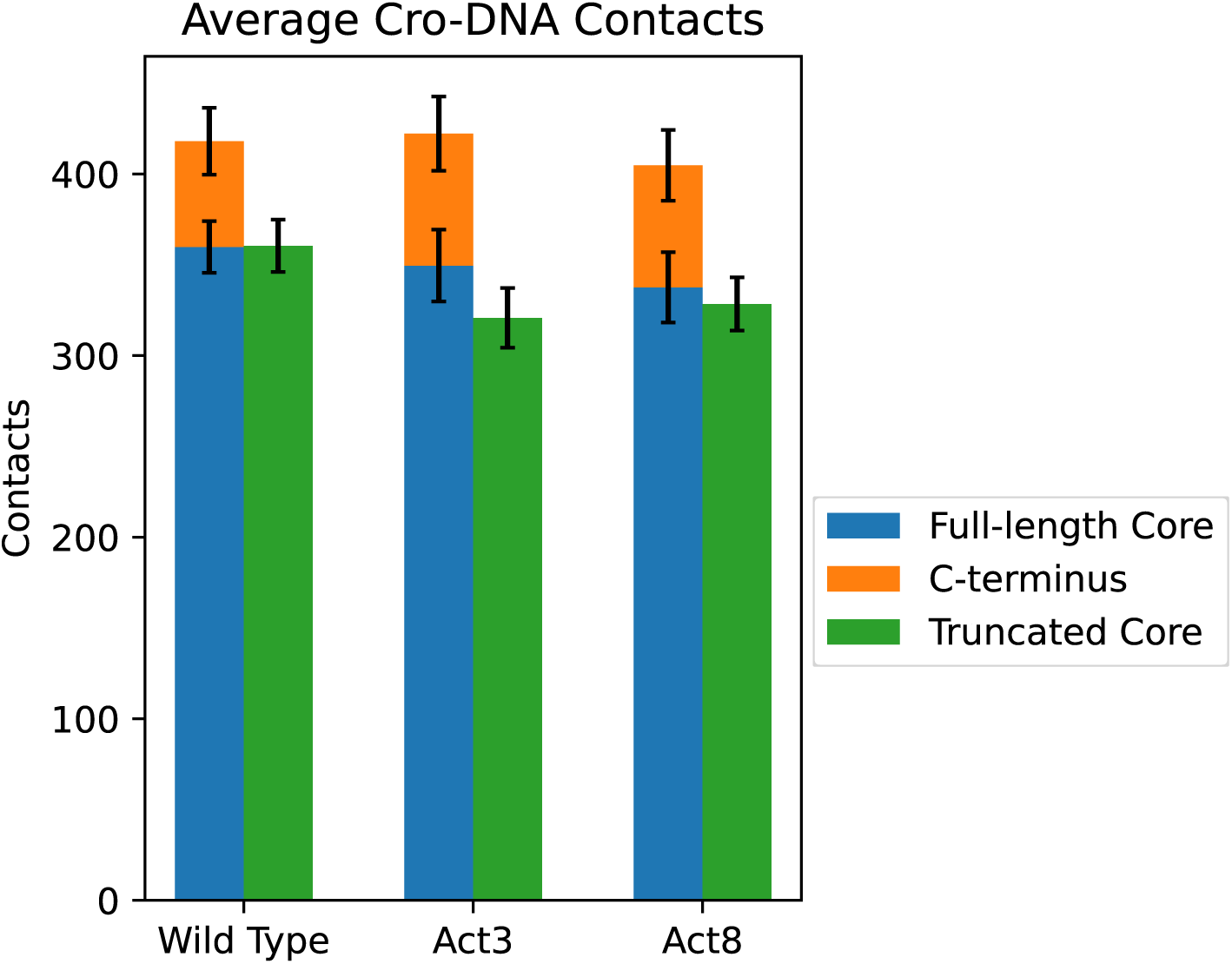
Average contacts between Cro subunits and DNA. In wild-type and Act8 variants, truncation of the C-terminal residues reduced DNA contacts only by the amount contributed by those residues. In Act3 variants, truncation also caused additional losses in core–DNA contacts beyond those attributable to the removed residues.

We used MM/GBSA analyses to further quantify the differences in Cro–DNA binding caused by C-terminal truncation. Only in the Act3 mutant did truncation reduce binding affinity: full-length Act3 Cro had an average binding energy of –138.8*±*6.2 kcal/mol, which became less favorable to –112.8*±*4.3 kcal/mol upon truncation (Table 2). Full-length wild-type Cro had –116.6*±*15.7 kcal/mol, compared to –118.5*±*7.5 kcal/mol for its truncated form, while full-length Act8 had –103*±*25.1 kcal/mol versus –119.4*±*8.3 kcal/mol when truncated. Notably, this analysis was performed only on frames in which both Cro subunits remained associated with DNA and does not account for truncated Act3 conformations after subunit dissociation. Together, the combined results from dissociation frequencies, contact analyses, and binding energy calculations show that in Act3, the mutation sites and the disordered C-terminus act together to maintain DNA binding, and that disrupting both significantly reduces this ability.

**Table 2:** MM/GBSA analysis of Cro-DNA binding affinity. Act3 mutations showed reduced affinity from truncation.

| Cro Variant | $\Delta E_{elec}$ | $\Delta E_{internal}$ | $\Delta E_{VdW}$ | $\Delta E_{total}$ |
| --- | --- | --- | --- | --- |
| Full-Length Wild Type | $-19.6 \pm 6.8$ | $15.7 \pm 9.4$ | $-152.0 \pm 10.6$ | $-116.6 \pm 15.7$ |
| Truncated Wild Type | $14.1 \pm 3.3$ | $12.3 \pm 5.6$ | $-144.9 \pm 3.9$ | $-118.5 \pm 7.5$ |
| Full-Length Act3 | $22.4 \pm 2.8$ | $11.3 \pm 4.5$ | $-172.6 \pm 3.3$ | $-138.8 \pm 6.2$ |
| Truncated Act3 | $27.0 \pm 1.9$ | $12.3 \pm 3.2$ | $-152.1 \pm 2.2$ | $-112.8 \pm 4.3$ |
| Full-Length Act8 | $24.3 \pm 10.7$ | $9.4 \pm 14.2$ | $-137.1 \pm 17.6$ | $-103 \pm 25.1$ |
| Truncated Act8 | $11.9 \pm 3.7$ | $9.5 \pm 6.2$ | $-140.8 \pm 4.2$ | $-119.4 \pm 8.3$ |

### 3.4 Cro Mutations Redistribute Contacts and Allosteric Coupling

Having characterized local changes from mutations and the behavior of the C-terminal tails, we next asked whether point mutations in Act3 and Act8 reorganize Cro’s global structure and dynamic coupling. In particular, we examined how mutations shift intra– and inter-subunit interactions in DNA-free and DNA-bound states, and whether these shifts relate to the DNA-binding differences described earlier.

Across all variants, full contact maps revealed the same core structure: residues 1–6 pair with residues 41–45 as part of a small β-sheet that also includes residues 49–57, while residues 7–36 form a three-helix bundle connected by turns. Each helix interacts with parts of the β-sheet, and in dimers the third β-strand from each subunit pairs to form a combined sheet. DNA-bound Cro maintains these internal contacts while adding extensive DNA interactions, primarily through helix 3 and, to a lesser extent, helix 2. This conserved core architecture across wild type and mutant forms, as well as in both free and DNA-bound states, provides a stable baseline from which mutation-driven changes can be identified.

Although this core is preserved, mutations introduce subtle shifts in contact patterns. To quantify these shifts, we first compared difference contact maps between wild type and each mutant (Figures S12–S13). This revealed several shared features: both mutations showed increased contacts at the dimer interface, reduced contacts between helix 2 and the C-terminal tail, and stronger interactions between the strand–turn–strand portion of the β-sheet and a mutation-adjacent region of helix 2. Act8 dimers also showed tighter association between the N-terminus and the β-sheet compared to Act3. When the dimers were bound to DNA, these differences persisted and, in some cases, became more pronounced, particularly at the protein–DNA interface (Figure 7). Some of the differences observed in dimers persisted in the DNA-bound context, while others, such as the N-terminal β-sheet association in Act8, became less prominent. Watson–Crick pairing of the DNA strands appeared slightly stronger (by 3–8%) in Act3 simulations, which may be a consequence of our setup: Act3–DNA complexes were initiated from equilibrated wild type–DNA frames. This difference could therefore reflect either a protocol artifact or a cooperative effect of altered Cro–DNA interactions described earlier.

**Figure 7:**
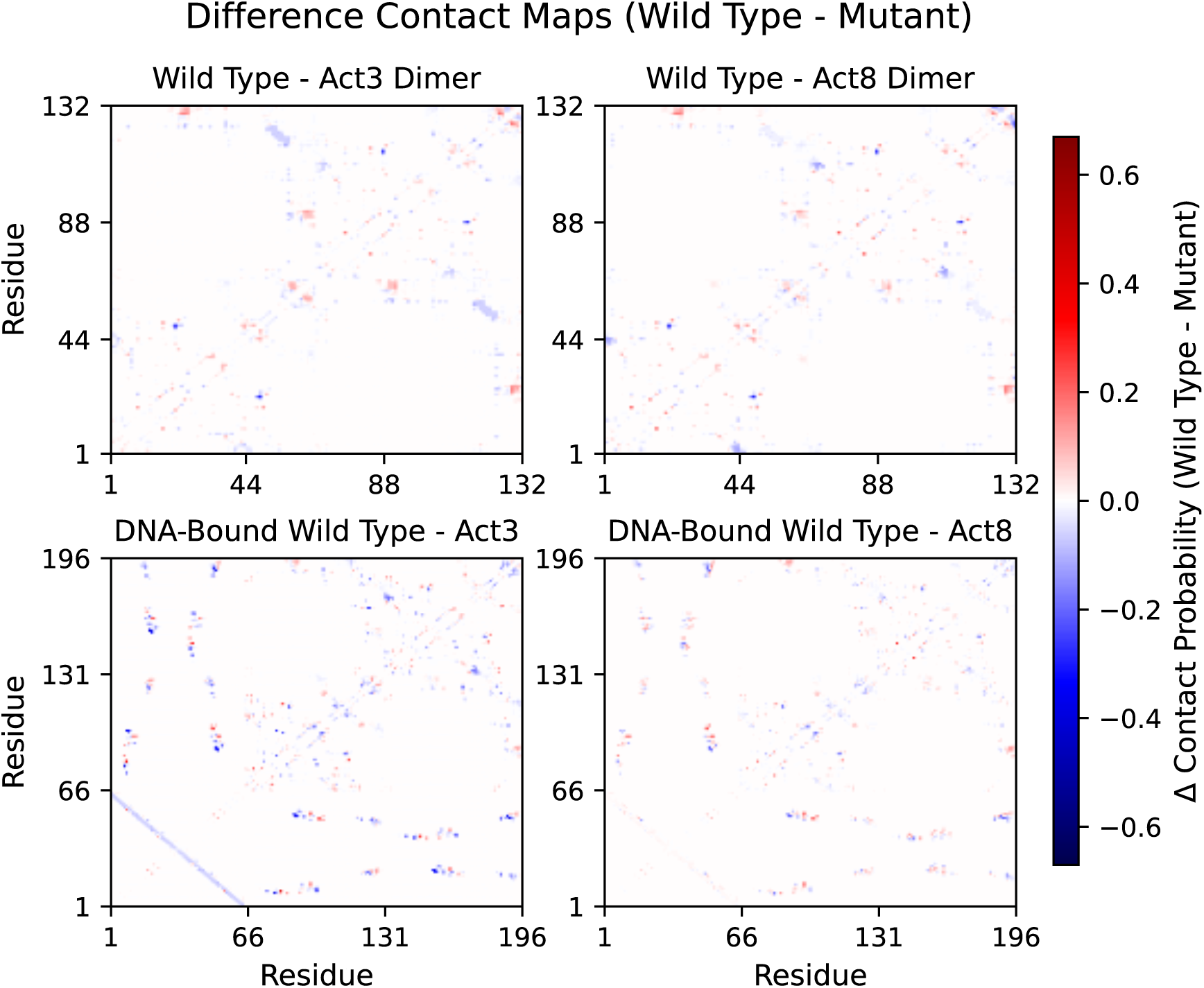
Difference contact maps between dimers (top) and DNA-bound dimers (bottom). Red regions indicate contact losses upon mutation and blue regions indicate contact formations upon mutations.

To better visualize and interpret these contact differences, we applied difference contact network analysis (dCNA) to full-length Cro dimers in both DNA-free and DNA-bound contexts. This method compares residue–residue contact probabilities between systems and groups residues into communities that are more internally connected than externally, allowing us to track shifts in coupling between these communities across systems.

In DNA-free systems, both Act3 and Act8 showed similar deviations from wild type, with no substantial divergence between the two mutants or between subunits within each simulation. Several shared features emerged. Mutations weakened coupling between helices within each subunit, including reduced contacts between helix 2 and helix 3 (e.g., community 3 to 4 in subunit 1 and 9 to 10 in subunit 2), as well as between helix 1 and helix 2 (communities 2 and 3 in subunit 1 and 8 and 9 in subunit 2) (Figure 8C–D). While modest in magnitude (typically 0.3–0.4), these reductions were consistent across variants and subunits. These intramolecular losses were accompanied by stronger coupling between subunits, particularly involving the C-terminal β-strand region and its interactions with the opposing subunit. For example, community 12 (C-terminal of subunit 2) formed new contacts with community 2 (helix 1 of subunit 1), and community 6 (C-terminal of subunit 1) with community 8 (helix 1 of subunit 2). Inter-subunit coupling between communities 2 and 12, 6 and 8, and 2 and 8 increased by 1.0 to 1.9 in the mutants relative to wild type. This shift suggests that when stabilizing interactions within a subunit are weakened, residues at the dimer interface, particularly those near the C-terminal strand, become more engaged, potentially compensating for the internal loss and reshaping dimer architecture in the absence of DNA.

**Figure 8:**
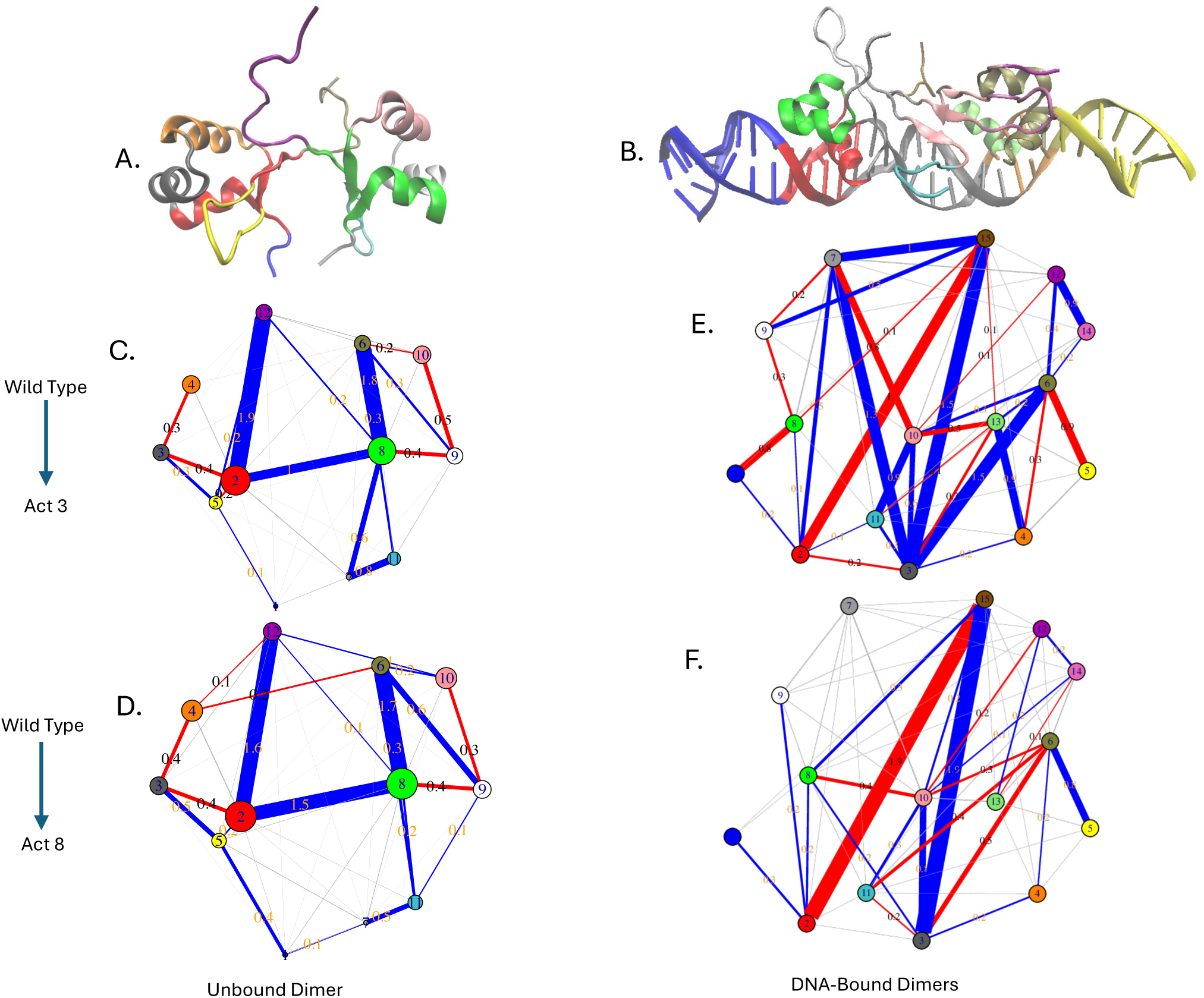
Difference contact network analysis (dCNA) results. (A) The Cro dimer was divided into 12 communities corresponding to secondary structure elements and their associations, including each subunit’s β-sheet. This community mapping was maintained when comparing wild-type Cro to Act3 (C) and Act8 (D). (B) Cro dimers bound to 64 base pairs of DNA were divided into 15 communities representing distinct interaction regions. Similar comparisons were performed for wild type to Act3 (E) and wild type to Act8 (F) variants.

In contrast to the DNA-free state, the DNA-bound dimers showed clearer mutation-specific differences. Community 3 (DNA between the binding sites and the dimer interface) formed more contacts with the communities comprising Act3’s β-sheet. Mutation-adjacent regions in both subunits (communities 6 and 8) showed reduced contacts with end DNA in Act3, but increased contacts in Act8 simulations (communities 1 and 5). Community 2, comprising the DNA-binding helix and the four base pairs it contacts most tightly in Cro subunit 1, had reduced interactions with the C-terminal residues of Cro subunit 2 (community 15) in both mutants. In these cases, subunit 2’s C-terminus tended to interact more with DNA between the Cro subunits, rather than with DNA adjacent to the DNA-binding helix and mutation site (Figure 8E–F).

Taken together, these results indicate that while the core Cro structure is maintained, mutations subtly redistribute contacts within and between subunits. In the absence of DNA, this redistribution enhances inter-subunit coupling near the C-terminal β-strand, potentially compensating for weakened intramolecular contacts. In the DNA-bound state, mutation-specific differences emerge at the protein–DNA interface, suggesting that Act3 and Act8 alter the dynamic balance between intra-protein stabilization and direct DNA engagement in distinct ways.

## 4 Discussion

In this study, we used all-atom MD simulations to investigate how engineered gain-of-function mutations and C-terminal truncation reshape the structure, dynamics, and functional potential of the λ Cro transcription factor. Cro normally functions as a DNA-bound dimer that represses transcription by blocking RNA polymerase binding.^55^ A small set of engineered substitutions can convert Cro into a dual-function factor that also recruits RNA polymerase,^23^ but the structural basis for this transformation has remained unclear. Our simulations provide atomistic evidence for mutation-induced surface remodeling, altered hydrogen-bonding networks, and dynamic rearrangements that influence both DNA contacts and accessibility of a distinct surface patch.

Most of the gain-of-function mutations in Act3 and Act8 cluster within residues 17–22 in helix 2, a segment that is largely solvent-exposed in the DNA-bound state but still makes frequent DNA contacts and hydrogen bonds in our simulations. These substitutions introduce substantial chemical and electrostatic changes: in Act3, residue 17 changes from a polar threonine to a hydrophobic valine, while residue 21 shifts from a positively charged lysine to a nonpolar alanine; in Act8, residue 17 remains threonine, but residue 21 is replaced by valine, reducing polarity and removing positive charge. Together, these mutations alter local hydrophobicity, polarity, and surface electrostatics, creating a more hydrophobic surface patch that is positioned and chemically suited for interaction with the N-terminal domain of the RNA polymerase σ subunit.^56,57^ The mutated variants also showed reduced contact between the C-terminal tail and this region, suggesting a cooperative rearrangement that preserves patch accessibility for potential binding partners. Outside the patch, residue 26 showed increased burial and hydrogen bonding in both Act3 and Act8, behavior that may partially compensate for reduced DNA contacts from residue 17. Because Cro–DNA association often begins with nonspecific recruitment before scanning to the preferred operator sequence,^58–60^ these compensatory changes may help preserve recruitment efficiency even if final sequence-specific binding is altered.

The disordered C-terminal tail plays a context-dependent role in DNA interactions and is experimentally required for Act3 function. This region contains lysines at positions 62 and 63, which often stabilize DNA contacts in other systems through electrostatic interactions.^61^ In our simulations, the tail frequently contacted DNA in all variants, yet MM/GBSA assigned little energetic weight to its presence, which is likely due to shielding by nearby polar residues or underestimation of electrostatics in implicit solvent models. Only truncated Act3 showed a significant drop in DNA affinity and loss of association, matching experimental observations that truncation eliminates its dual function.^23^ Because the C-terminus is thought to support nonspecific DNA binding,^16,62^ loss of these lysines alone may be sufficient to suppress recruitment. However, the disassociation of truncated Act3 suggests that a combination of nonspecific and sequence-specific interactions is essential for function, and that tail loss may synergize with mutation-induced surface remodeling to alter recruitment and binding behaviors.

MD simulations provide detailed molecular-scale views, yet they capture only part of the conformational space accessible to Cro. Our work began from three experimentally determined configurations that reflect local energy minima rather than the complete folding landscape. This starting point may cause truncated Cro dimers to appear more stable than if simulated from an unfolded state. We also did not simulate the full progression from monomer to dimer to DNA-bound complex, which limits direct assessment of how mutations and truncations influence assembly. Even with these constraints, our findings reveal clear mechanistic links between specific mutations, C-terminal dynamics, and altered DNA-binding behavior. By showing how minimal changes to sequence can reconfigure interaction surfaces and reshape cooperative binding, this work advances our understanding of how small transcription factors can be rationally redesigned. Such insight not only informs efforts to repurpose Cro but also offers a general framework for engineering compact regulators with customized functions in synthetic biology and gene control applications.

## 5 Data availability

All simulation inputs, scripts used for analysis, and processed data files are available at https://github.com/WereszczynskiGroup/supplemental-data-hebert-cro-2025. Raw trajectory data is available on Zenodo at https://doi.org/10.5281/zenodo.16877368. These trajectories are water-stripped and temporally strided to reduce file size but cover the full duration of each simulation. Together, these resources are sufficient to reproduce all analyses and key results reported in the manuscript.

## Acknowledgments

This project was supported by the National Institutes of Health grant R35GM119647. During the preparation of this work the authors used ChatGPT to refine the manuscript language. After using this, the authors reviewed and edited the content as needed and take full responsibility for the content of the publication.

