## Supplemental Material for "Molecular Mechanisms of Gain-of-Function Mutations in λ Cro Revealed by Molecular Dynamics Simulations"

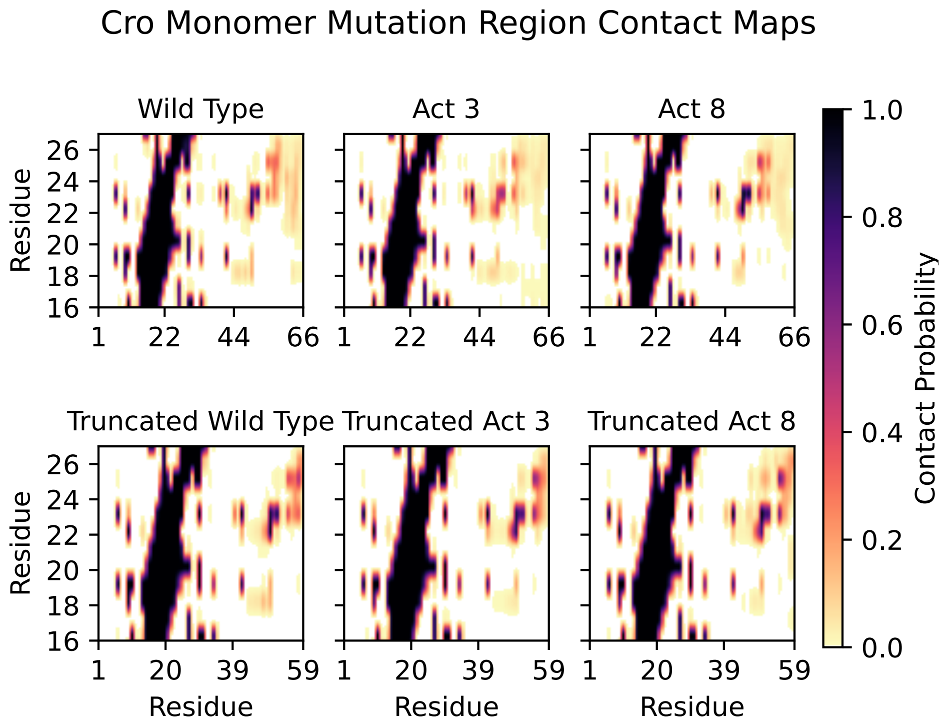


Figure S1: Contact maps of the mutation region of Cro monomers against the entire monomer.

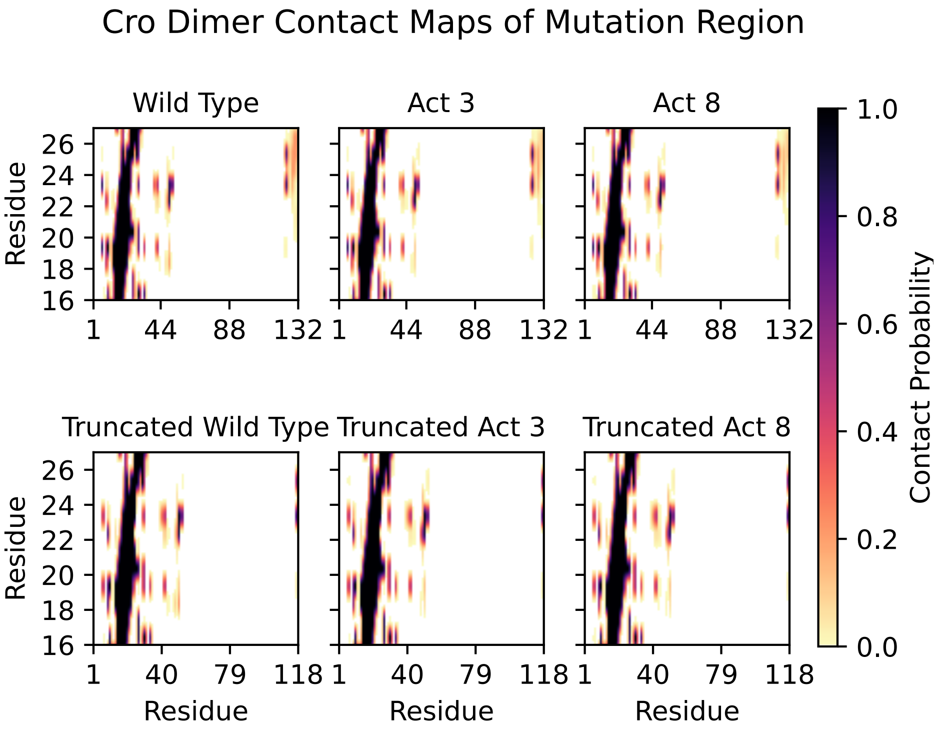


Figure S2: Contact maps of the mutation region of a subunit of a Cro dimer against the dimer complex.

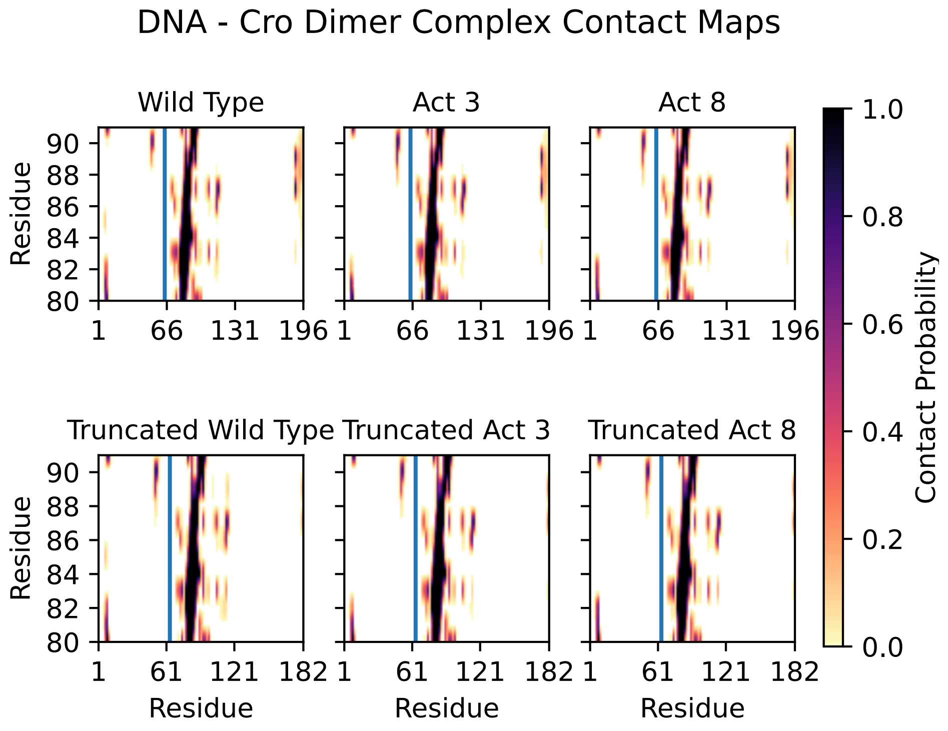


Figure S3: Contact maps of the mutation region of a subunit of a Cro dimer against the DNA-dimer complex. Residues 1-64 represent DNA.

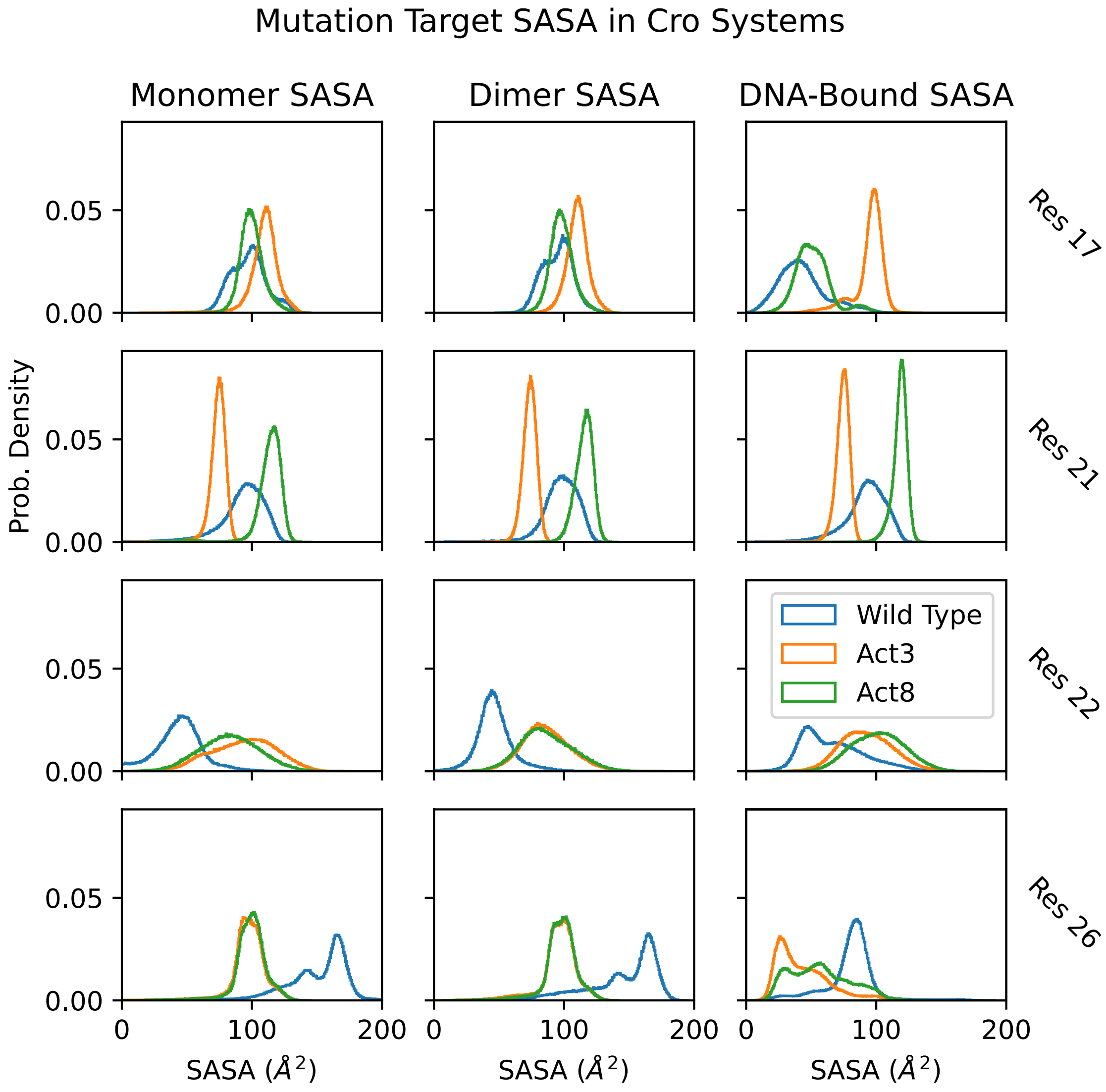


Figure S4: Solvent accessible surface area of mutation targets in Cro simulations. Differing amino acid geometries lead to changes in burial/accessibility.

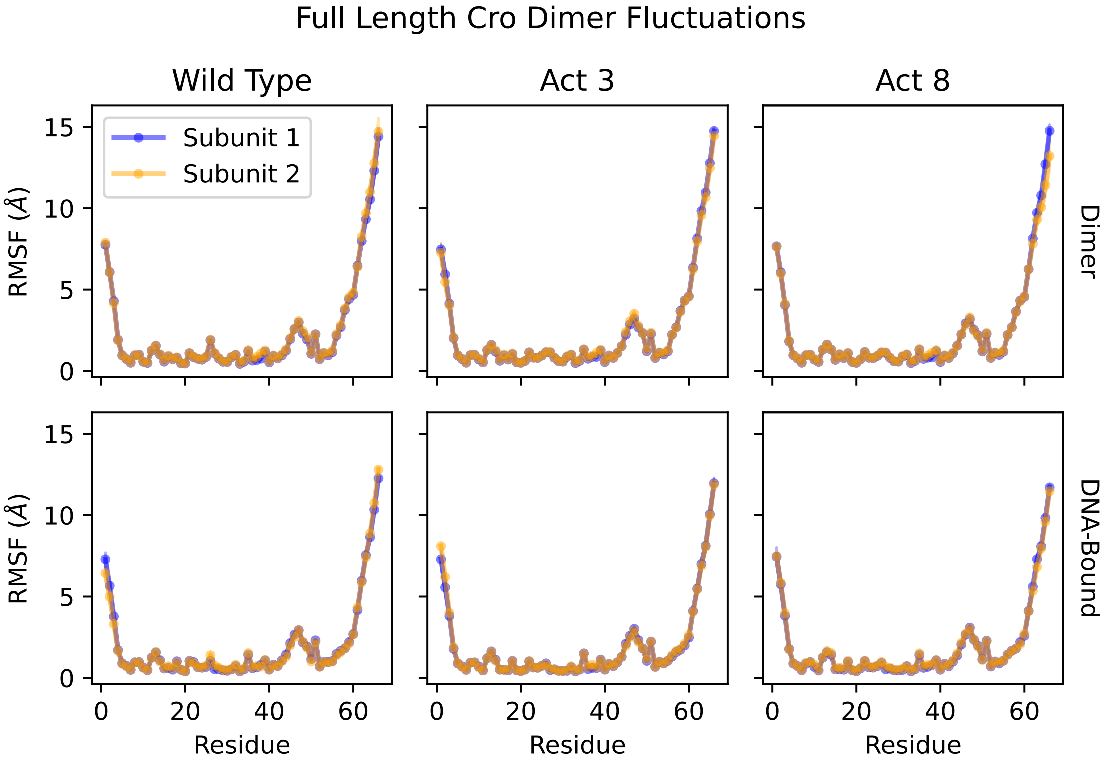


Figure S5: Root mean squared fluctuations of full-length, dimerized Cro subunits. Dimerized Cro subunits in similarly folded states experience symmetric fluctuations. Top row: Dimerized Cro. Bottom: DNA-bound Cro dimers.

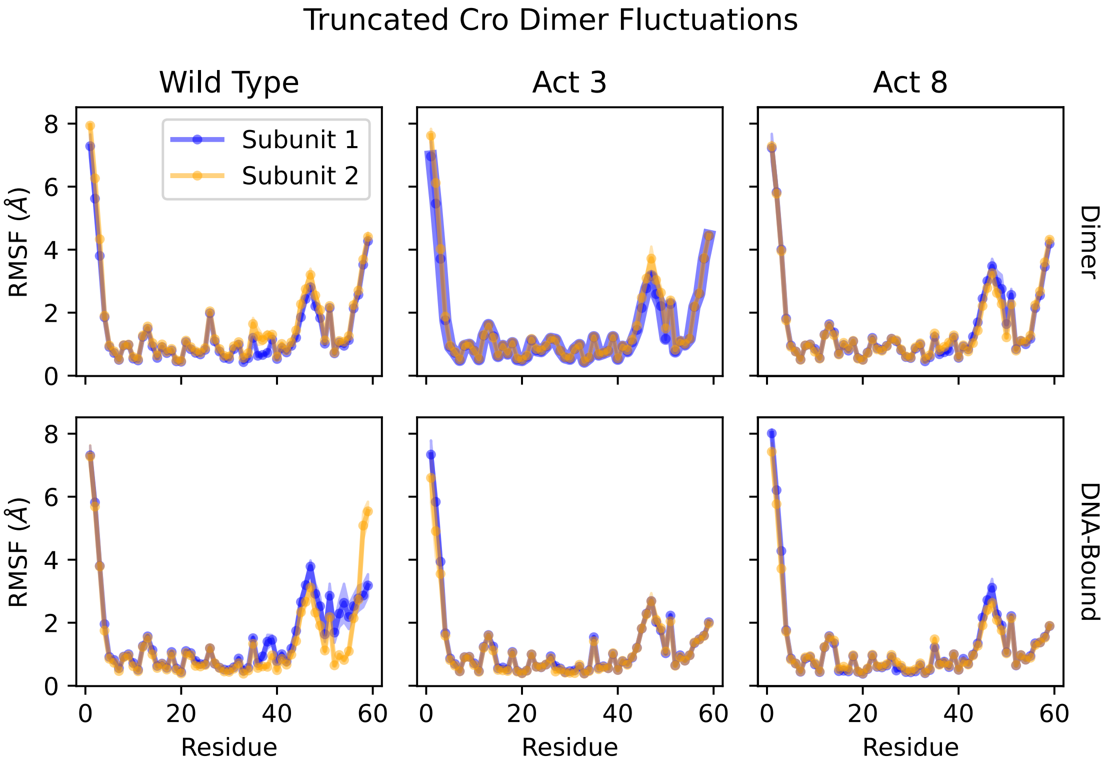


Figure S6: Root mean squared fluctuations of truncated, dimerized Cro subunits. Dimerized Cro subunits in similarly folded states experience symmetric fluctuations. Top row: Dimerized Cro. Bottom: DNA-bound Cro dimers.

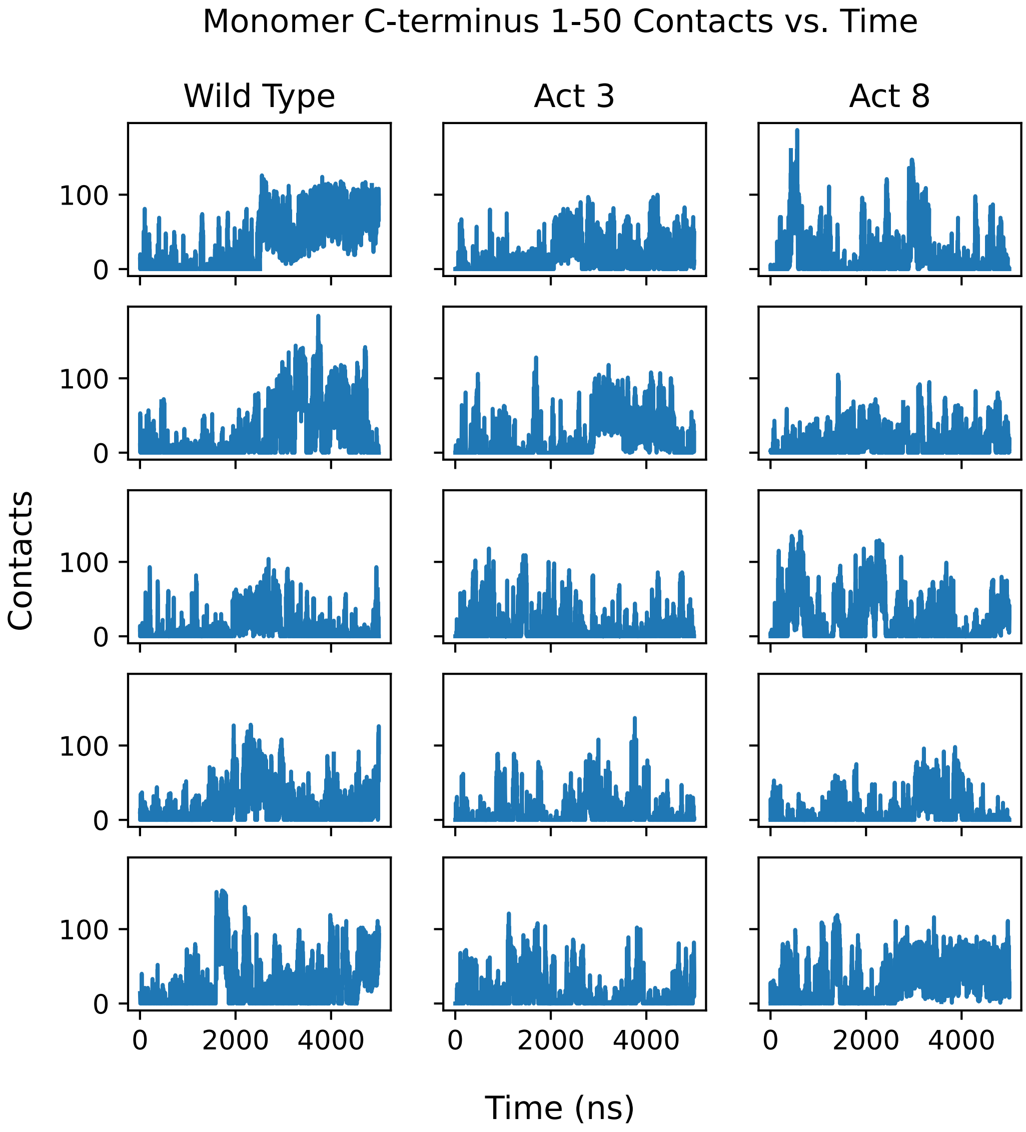


Figure S7: Contacts over time between residues 1-50 and residues 60-66 in full length Cro monomers. High variation within each run indicates transient contacts.

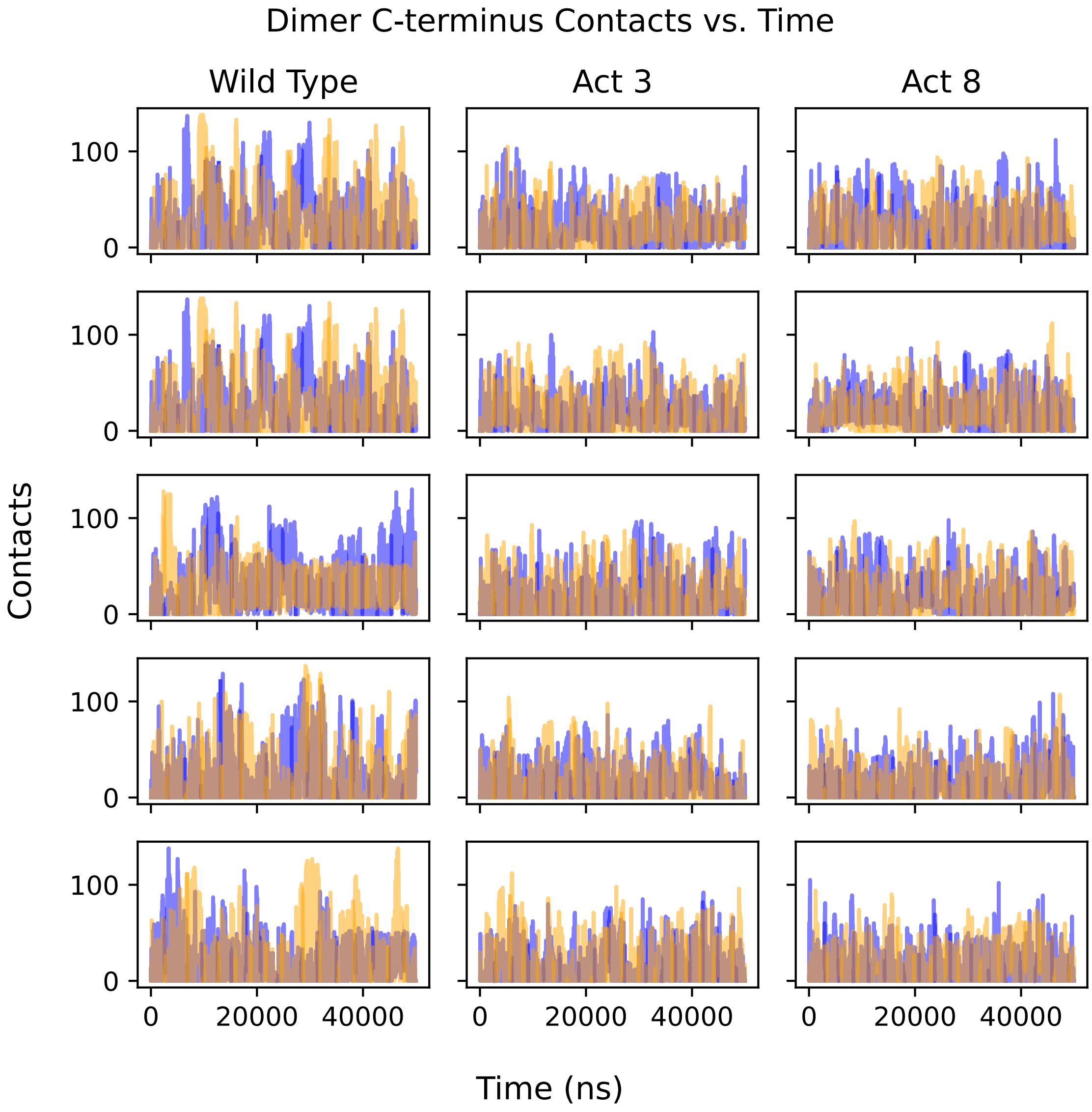


Figure S8: Contacts over time between residues 60-66 in each Cro subunit and residues 1-50 of both subunits in full length Cro dimers. High variation within each run indicates transient contacts.

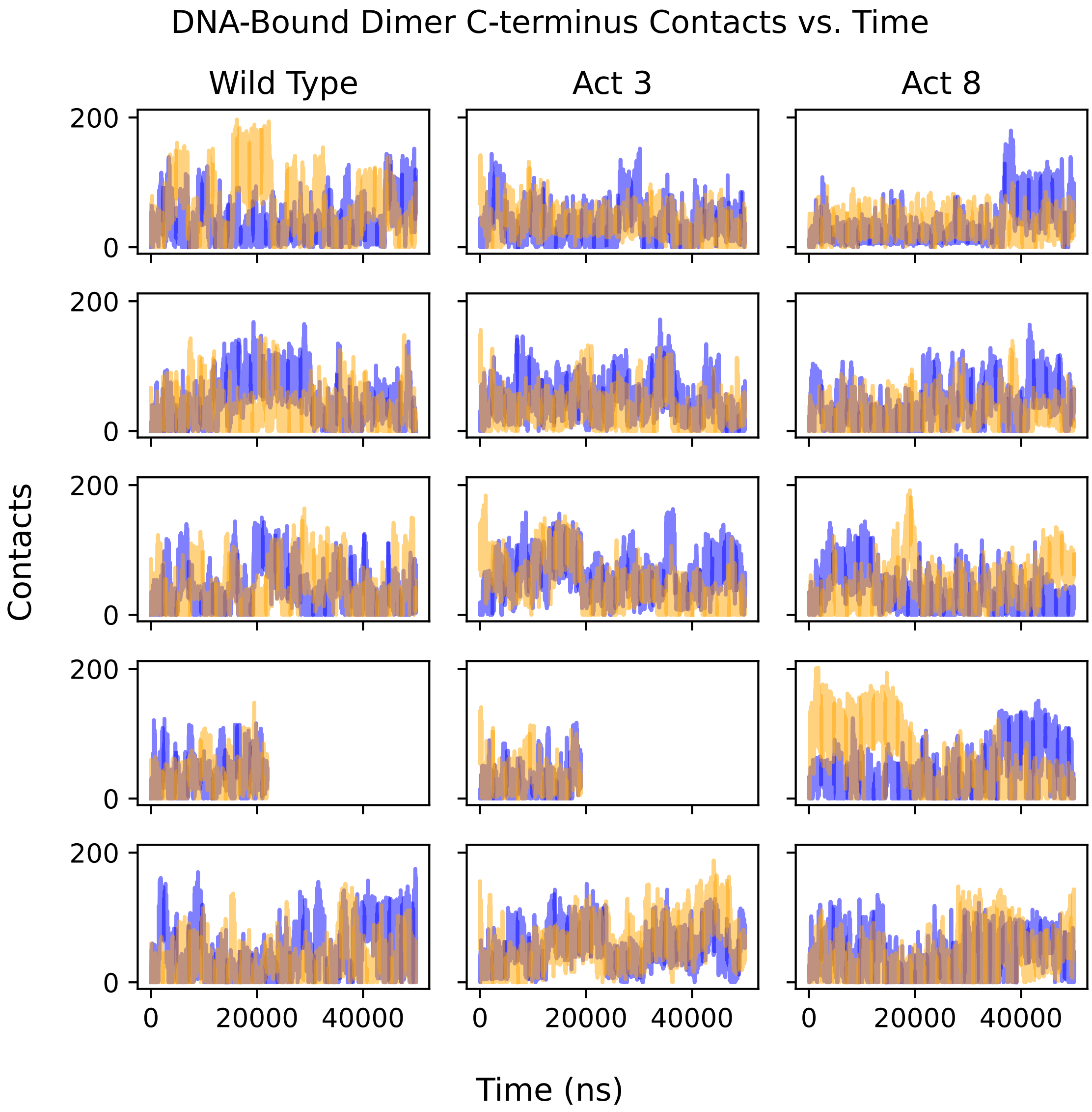


Figure S9: Contacts over time between residues 60-66 in each Cro subunit, DNA, and residues 1-50 of both subunits in full length Cro dimers. High variation within each run indicates transient contacts.

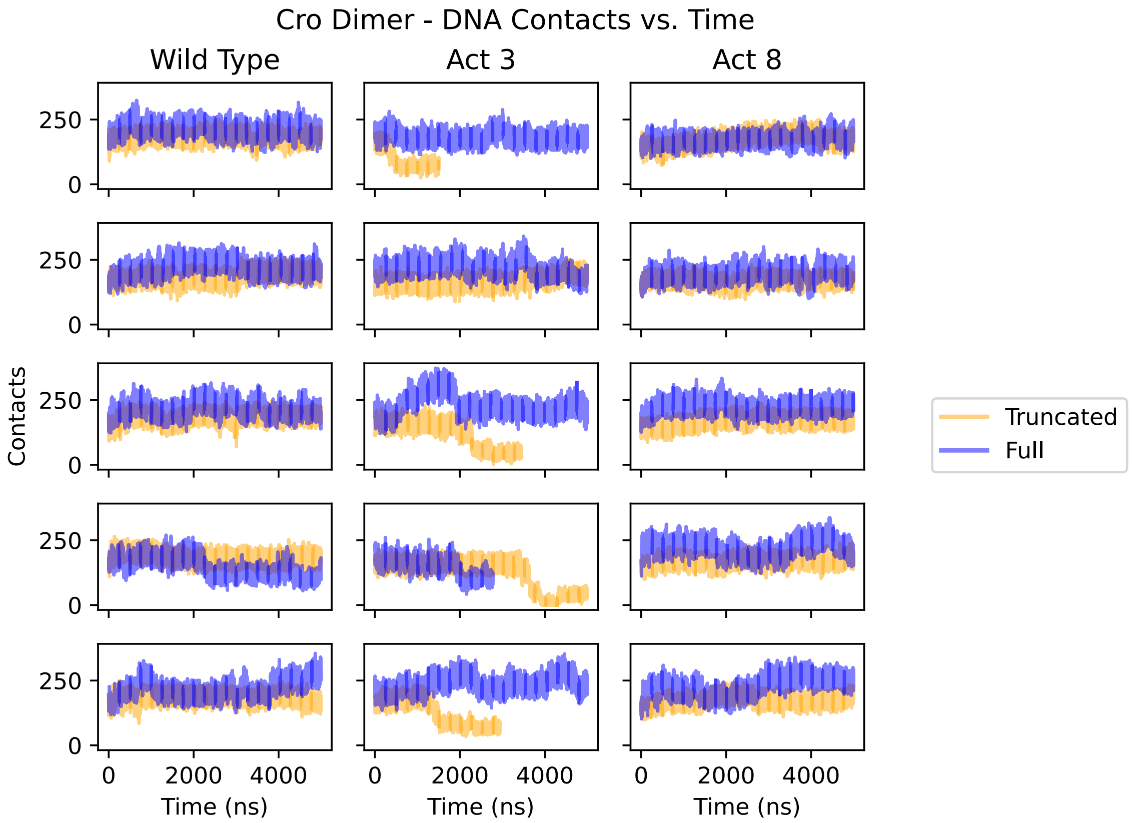


Figure S10: Contacts over time between both Cro subunits and DNA. Disassociation between one Cro subunit and DNA is associated with the drops seen in full length wild type's and Act3's fourth simulations, as well as the drops seen in all but the second simulation of truncated Act3. We did not continue most simulations after this disassociation.

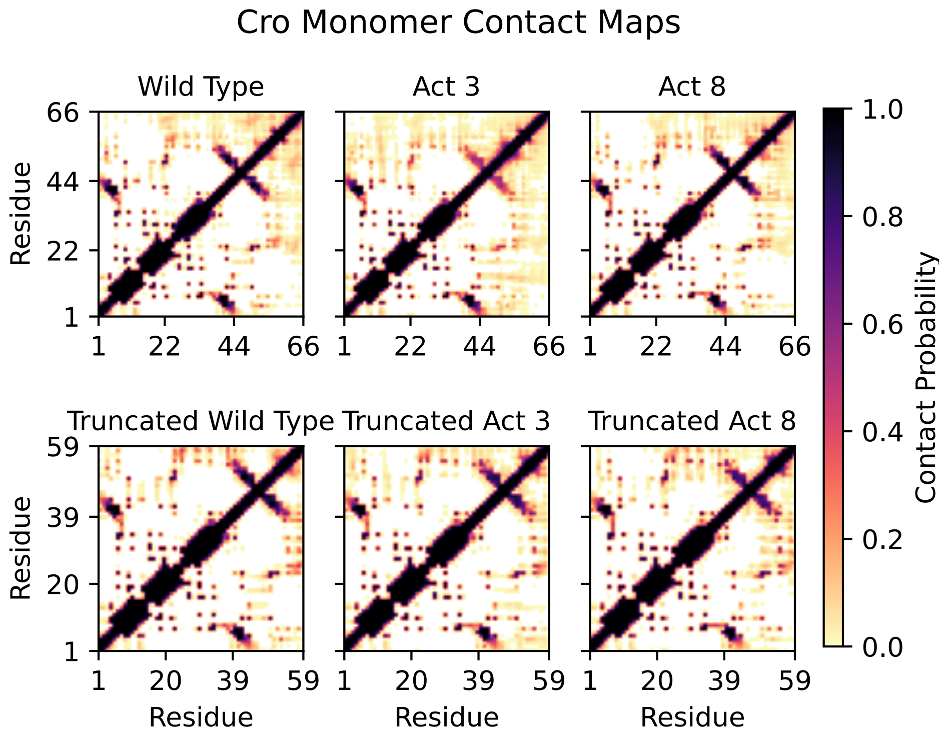


Figure S11: Contact maps of Cro monomer structures. The final βstrand is prone to unfolding, smearing out contacts from the C-terminus.

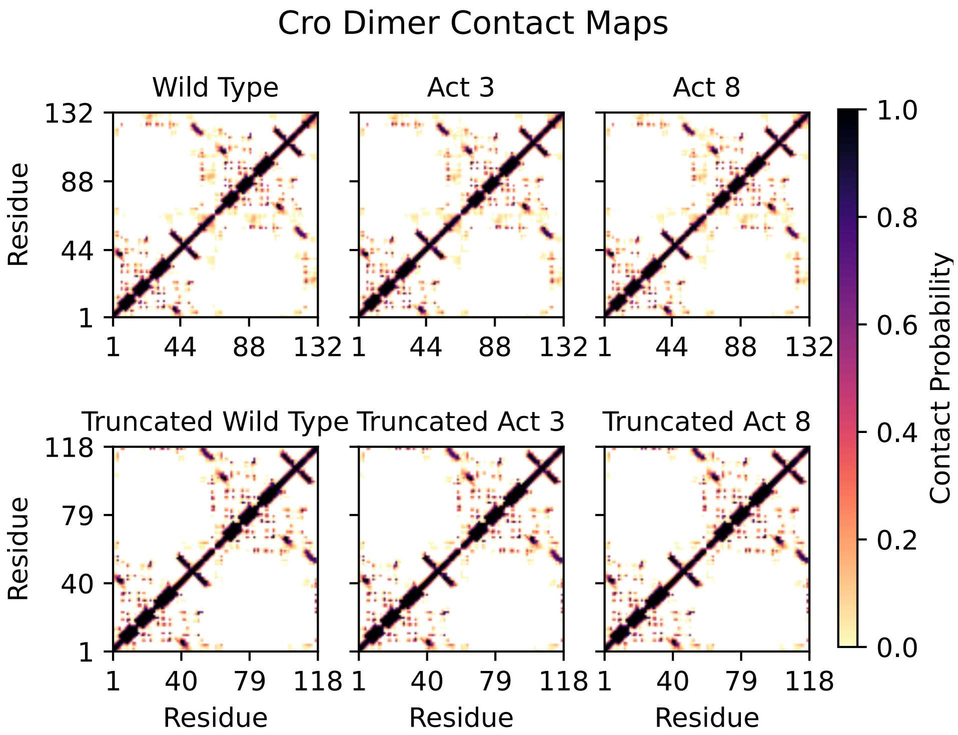


Figure S12: Contact maps of Cro dimer structures. Top: Residues 1-66 correspond to the first Cro subunit, and the remainder the second Cro subunit. Bottom: Residues 1-59 correspond to the first Cro subunit, and the remainder the second Cro subunit.

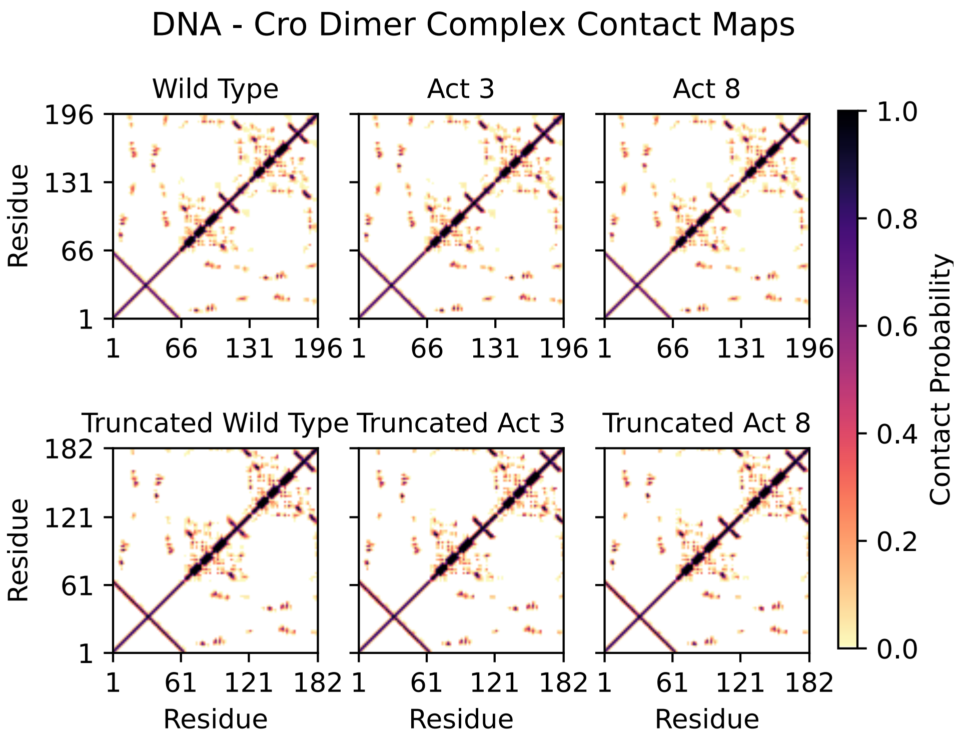


Figure S13: Contact maps of DNA-bound Cro dimer structures. Residues 1-64 represent DNA. Top: Residues 65-130 correspond to the first Cro subunit, and residues 131-196 correspond to the second Cro subunit. Bottom: Residues 65-123 correspond to the first Cro subunit, and residues 124-182 correspond to the second Cro subunit.
